# Anterior cingulate neurons display subregion-specific interaction with frontal eye fields as revealed by combined antidromic stimulation and resting state imaging

**DOI:** 10.1101/2024.11.09.622778

**Authors:** Sahand Babapoor-Farrokhran, Alex J. Major, Kevin D. Johnston, Earl K. Miller, Joan A. Camprodon, Stefan Everling

## Abstract

The anterior cingulate cortex (ACC) is thought to exert cognitive control over saccade generation in the frontal eye fields (FEF), but the nature of this interaction remains unclear. Although prior imaging studies have suggested ACC interacts with FEF, few studies have confirmed this by electrophysiological recordings. This study aimed to characterize the functional connectivity between ACC and medial and lateral FEF during cognitive saccade tasks. We combined resting-state functional MRI (rs-fMRI) with single-unit electrophysiology in two macaque monkeys performing memory-guided saccade and pro-/anti-saccade tasks. Anti- and ortho-dromic stimulation was used to electrophysiologically identify ACC neurons mono- and polysynaptically connected to FEF. We analyzed ACC neuronal selectivity for different task aspects and correlated these properties with both positive and negative rs-fMRI functional connectivity between ACC and FEF subregions. Anti- and ortho-dromically identified ACC neurons were predominantly connected to medial FEF, which showed stronger positive functional connectivity with ACC compared to lateral FEF. Sites with higher proportions of task-selective neurons yielded stronger functional connectivity with FEF. This stronger functional connectivity was particularly related to the delay and saccadic periods of different cognitive saccade tasks. Using combined imaging and electrophysiology, our findings provide converging evidence for functional interactions between ACC and FEF, predominantly medial FEF regions which encode large amplitude saccades. The correlation between functional connectivity and task-related neuronal selectivity supports ACC’s interaction with FEF in the modulation of saccade generation and cognitive control. Additionally, we report suggestive evidence that mono- and poly-synaptic connections may be related to positive functional connectivity, but we found no such relationship for negative functional connectivity (anticorrelations). These results advance our understanding of prefrontal cortical interactions in oculomotor control and the electrophysiological mechanisms of positive and negative resting-state functional connectivity.

## Introduction

A key component of cognitive flexibility is altering motor outputs in a context-dependent manner. A flashed light in the peripheral visual field will reliably produce an automatic saccadic eye movement towards this stimulus. However, suppressing the generation of this prepotent saccade in frontal eye fields (FEF) and downstream superior colliculus (SC) requires top-down influence from high-order cognitive regions. Anterior cingulate cortex (ACC) is a large and multifunctional brain region, well-positioned to provide influence over saccadic outputs of FEF. In turn, FEF likely provides bottom-up input to ACC (Babapoor-Farrokhran et al., 2017) possibly providing ACC with the context-related information. ACC neurons have been implicated in conflict monitoring between two conflicting stimuli or motor outputs (Johnston et al., 2007; Womelsdorf et al., 2010) and monitoring of performance and reward (Ito et al., 2003). The ACC is well-connected to incorporate both high-order cognitive processing (e.g., connections with other prefrontal areas) and emotional affect (through its connections with the motivation and reward circuits such as ventral tegmental area, (Devinsky et al., 1995)) in order to influence FEF’s control of saccadic outputs.

Tracer studies have confirmed direct anatomical projections between ACC and FEF (Barbas and Mesulam, 1981; Bates and Goldman-Rakic, 1993; Stanton et al., 1995; Wang et al., 2004). However, less is understood about the functional role of FEF-projecting ACC neurons’ spiking activity. Previous functional imaging studies have confirmed ACC activation during saccades in humans (Paus et al., 1993; Petit et al., 1993; Picard and Strick, 1996; Gaymard et al., 1998; Raemaekers et al., 2005; Brown et al., 2006; Amiez and Petrides, 2014) and have demonstrated functional connectivity between ACC and FEF in macaques (Hutchison and Everling, 2012; Babapoor-Farrokhran et al., 2013). Indeed, both ACC and FEF are part of the dorsal attention network to coordinate cognitive behaviors (Kaping et al., 2011; Farrant and Uddin, 2015; Womelsdorf and Everling, 2015).

What is the nature of ACC’s influence over FEF or vice versa? Does ACC interact differently with medial which encodes large saccades versus lateral FEF which encode small amplitude saccades? We hypothesized that ACC neurons exert cognitive control over FEF, enhancing directional selectivity of saccadic targets and aiding in suppression of prepotent automatic saccades. Specifically, we investigated the functional connections between ACC (Brodmann area 24, also referred to as medial cingulate cortex (Saleem and Logothetis, 2012; Procyk et al., 2016)) and medial or lateral FEF using a combination of resting state functional magnetic resonance imaging (rs-fMRI) and electrophysiology during saccadic cognitive tasks. This region of ACC could be analogous to “cingulate eye fields” reported elsewhere (Wang et al., 2004). Here, we recorded from ACC neurons and identified those neurons by anti- and orthodromic microstimulation that were mono- and polysynaptically connected with FEF. Using this combination of functional imaging and advanced electrophysiology, our study functionally probed the proposed role of FEF-projecting ACC neurons and the electrophysiological underpinnings of positive vs negative rs-fMRI functional connectivity.

## Materials and Methods

Experiments were performed on two adult male naive macaque monkeys (one *Macaca mulatta*, weight 9 kg; one *Macaca fascicularis*, weight 7 kg). In each monkey, implantation of the recording chambers was performed over the right arcuate sulcus and right anterior cingulate sulcus based on MRI guidance. Further details for the surgical procedures have been described previously (Phillips and Everling, 2012). To confirm the location of the recording chambers and reconstruction of the recording sites, post-surgical MRIs were obtained. All experimental methods and procedures were carried out in accordance with the guidelines of the Canadian Council of Animal Care policy on the care and use of laboratory animals and approved by the Animal Use Subcommittee of the University of Western Ontario Council on Animal Care. The monkeys were closely monitored by the university veterinarians.

### Behavioral tasks

Animals were trained to perform a standard memory-guided saccade (Fig. 1A). The details of this task have been described previously (Babapoor-Farrokhran et al., 2017). Briefly, a white central fixation point (0.15°) presented is presented. Once a subject fixated on this spot for 500 ms, a white target stimulus (0.15°) was presented for 100 ms in one of eight directions with equal probability (0°, 45°, 90°, 135°, 180°, 225°, 270°, and 315°; 9° eccentricity). The animal was required to maintain central fixation during the stimulus presentation and during the 1000 ms delay epoch following stimulus presentation. During the delay epoch, the target stimulus was not visible, but the central fixation point remained illuminated. The disappearance of the central fixation point was the cue for the monkeys to perform a saccade toward the remembered target stimulus location. The monkeys were required to perform a saccade within 500 ms after the offset of the central fixation point and to maintain fixation at the remembered target location for a random duration of 300–600 ms to obtain a water reward. Upon the correct performance of all the required steps, the target stimulus reappeared and a water reward was delivered (Fig. 1A).

**Figure 1.**
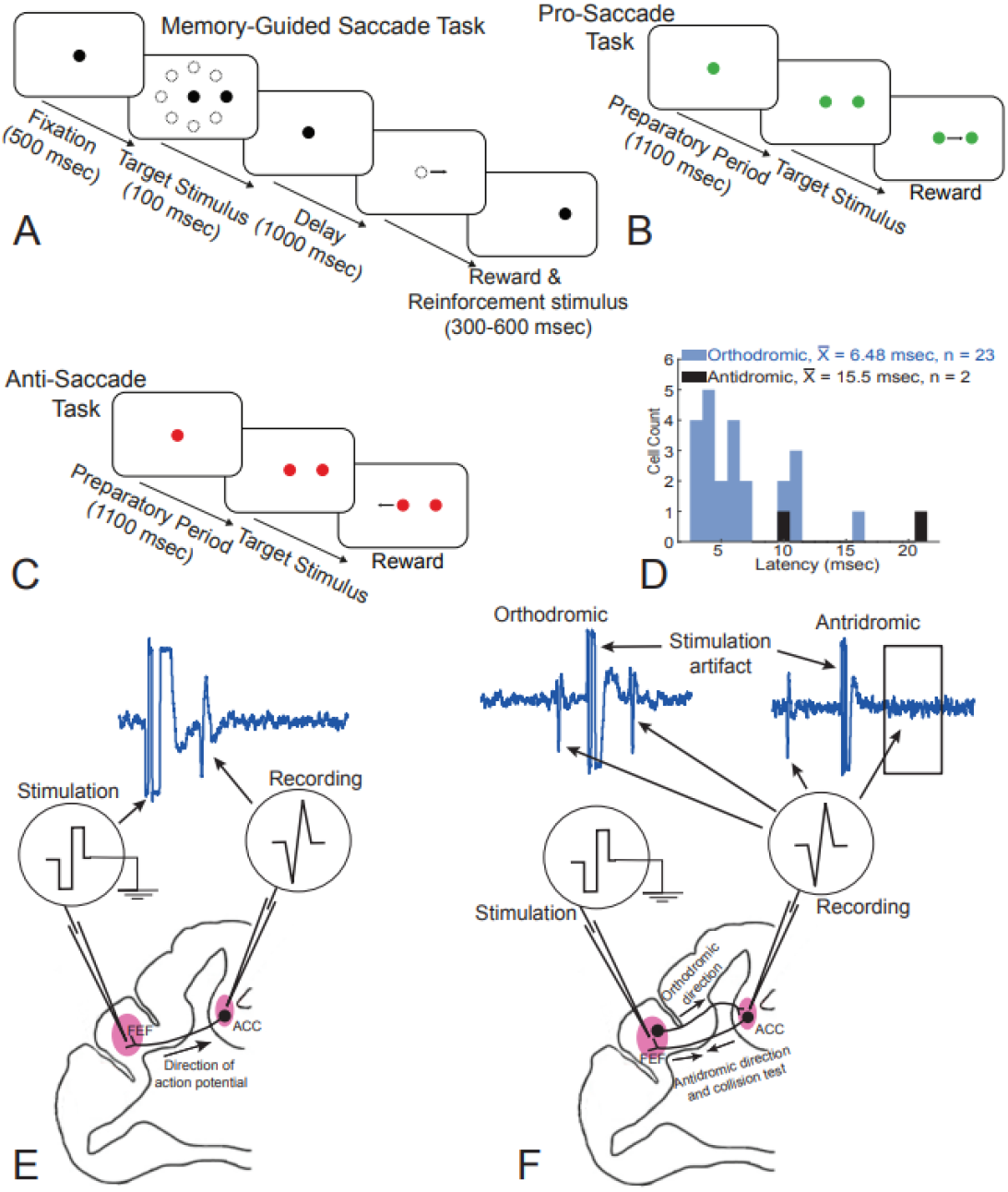
Behavioral tasks and microstimulation paradigm. Schematic of (A) memory-guided saccade task, (B) pro-saccade task, and (C) anti-saccade task. (D) Latency histogram of eliciting an action potential in anti- or orthodromically-identified ACC neurons following the stimulation in the FEF. (E) Schematic of a stimulated ACC neuron sending a projection to FEF. (F) Schematic of the collision test. An orthodromic neuron will fire an action potential following collision test whereas an antidromic neuron will not fire an action potential following the collision test.

The number of error trials in the memory-guided saccade task was low and this made the analysis of performance-related effects difficult. Therefore, we also utilized a randomly interleaved pro-/anti-saccade task. In this task, animal performance was lower, especially during anti-saccade trials. In this task, a central fixation point (0.15°) appeared with a green or red color to cue monkeys to perform either a pro- or anti-saccade, respectively. The fixation point remained illuminated for a period of 1100–1400 ms, which we refer to as the preparatory epoch. Monkeys were required to maintain fixation during the preparatory epoch. Next, a stimulus appeared (9° eccentricity) either to the right or left of the central fixation point. In pro-saccade trials, the monkeys made a saccade towards the target stimulus, and in anti-saccade trials, they were required to saccade in the opposite direction, away from the target stimulus (Fig. 5). The monkeys were required to maintain fixation at the target location for a random duration of 300– 600 ms to obtain a water reward.

Eye movements were recorded at 500 and 1,000 Hz using video eye trackers (EyeLink II and EyeLink 1000; SR Research, Ottawa, ON, Canada). Presentation of the behavioral stimuli, monitoring of the responses, and reward delivery were controlled using CORTEX experimental control software (NIMH, Bethesda, MD, USA). Task events, vertical and horizontal eye positions, and digitized neural signals were stored together using either a Plexon MAP or Omniplex System (Plexon Inc., Dallas, TX). Upon visual inspection of the eye traces, trials in which the eye movement trajectory deviated from the normal pathway were excluded.

### Rs-fMRI data acquisition

We utilized the same data acquisition procedures to obtain functional connectivity (FC) maps as utilized in previous studies (Hutchison et al., 2011, 2012; Babapoor-Farrokhran et al., 2013). The monkeys were anesthetized on the day of the procedure by intramuscular injections of ketamine hydrochloride (7.5 mg/kg), ipratropium (0.025 mg/kg), and atropine (0.4 mg/kg). Thereafter, we administered 3 mL of propofol (10 mg/mL) intravenously followed by oral intubation. Anesthesia was maintained with inhaled 1.5% isoflurane mixed with oxygen. Animals had spontaneous respiration during the image acquisition. A custom-built chair was used to position and head-fix the animal while in the magnet bore. At the time of acquiring functional images, isoflurane level was reduced to 1%. The following parameters were monitored during the image acquisition: end-tidal CO_2_ via capnometer (Covidien-Nellcor, Boulder, CO), rectal temperature via fiber-optic temperature probe (FISO, Quebec City, QC, Canada), and respiration via respiratory bellows (Siemens, Union, NJ). Vital signs were in the normal physiologic limits throughout the image acquisition (respiration rate: 24–32 breaths/min, temperature: 36.8°C, end-tidal CO_2_: 31–40 mmHg). Thermal insulation and a heating disk (SnuggleSafe, Littlehampton, West Sussex, UK) were used to maintain body temperature. Image acquisition was performed under general anesthesia because there are no training requirements, there are fewer motion artifacts, and less physiological stress. Functional connectivity and correlated blood oxygen level-dependent (BOLD) fluctuations under isoflurane anesthesia have been robustly maintained despite the vasodilator properties of isoflurane and its effects on cerebrovascular activity (Farber et al., 1997; Vincent et al., 2007; Hutchison et al., 2012). Functional and structural images were acquired using an actively shielded 7-Tesla 68-cm horizontal bore scanner with a DirectDrive console (Agilent, Santa Clara, CA) and a Siemens AC84 gradient subsystem (Erlangen, Germany) operating at a slew rate of 350 mT·m^-1^·s^-1^. A conformal five-channel transceive primate head radiofrequency coil (designed and manufactured in-house) was used for magnetic resonance (MR) image acquisition. The coil included an array of elements wrapped 270° circumferentially around the head. An automated three-dimensional mapping process over the imaging volume of interest was used for magnetic field optimization (B0 shimming). Each monkey had 10 runs of 150 continuous echo-planar imaging (EPI) functional volumes (TR = 2000 ms, TE = 16 ms, flip angle = 70°, matrix = 96×96, field of view [FOV] = 96×96 mm, acquisition voxel size 1×1×1 mm), each with a 5 min scan duration. Phase encoding in the left-right direction using GRAPPA at an acceleration factor of 2 was utilized to acquire EPI images. Navigator echo correction was used to correct for physiological fluctuations. Anatomic MR image was acquired along the same orientation as the functional images with a high-resolution gradient echo (GRE; TR = 1100 ms, TE = 8 ms, matrix = 256×256, FOV = 96×96 mm, acquisition voxel size = 375 μm × 375 μm × 1 mm). For every monkey, a T1-weighted anatomic image was also acquired (TE = 2.5 ms, TR = 2300 ms, TI = 800 ms, FOV = 96×96 mm, 750-μm isotropic resolution).

### Image preprocessing

FSL software was used to preprocess functional MRI data (http://www.fmrib.ox.ac.uk). This procedure included motion correction (6-parameter affine transformation), brain extraction, spatial smoothing (full-width at half-maximum = 2 mm), high-pass temporal filtering (Gaussian-weighted least-squares straight line fitting with σ = 100 s), low-pass temporal filtering (half-width at half-maximum = 2.8 s, Gaussian filter), and non-linear registration (nonlinear image registration tool: FNIRT, http://www.fmrib.ox.ac.uk/fsl/fslwiki/FNIRT) to the individual monkey’s T2-weighted image(Babapoor-Farrokhran et al., 2013). Global mean signal was not regressed out from the data due to its propensity for introducing extra anticorrelations (Murphy et al., 2009) and the possibility of removing physiologically important signals (Schölvinck et al., 2010).

### Statistical analysis of fMRI data

For each monkey, spherical seed regions (2 mm diameter) were selected individually in the medial and lateral FEF of the right hemisphere in the T2-weighted anatomic images (Fig. 5). The extracted mean time course of the signal for each seed region, subject, scanning session was used as the regressor in a generalized linear regression analysis. The regression model included the seed time series as predictors and also nuisance covariates (6 motion parameters, white matter, and cerebrospinal fluid [CSF]). White matter and CSF nuisance covariates were included in the model to further regress out the physiological noise from the data because cardiac and respiratory activity was monitored but not recorded during MR image acquisition. A similar method has been previously implemented to remove physiological noise in human and monkey resting-state fMRI (rs-fMRI) studies (Lund and Hanson, 2001; Margulies et al., 2007; Shim et al., 2010; Hutchison et al., 2011, 2012; Babapoor-Farrokhran et al., 2013). For further information about noise reduction of fMRI data, please refer to (Shmueli et al., 2007; Babapoor-Farrokhran et al., 2013).

The first level of the analysis was carried out on the individual subjects at each scanning session. In the second level, the analysis results from the first level used to generate functional connectivity maps across 10 scans for each monkey by implementing a second-level fixed-effects analysis using the FSL software package (http://www.fmrib.ox.ac.uk). These functional connectivity maps guided us to place recording chambers over ACC regions with functional connectivity to FEF.

### Electrophysiological Recording and Microstimulation Protocol

In electrophysiological experiments, three dura-puncturing tungsten microelectrodes (FHC, Bowdoinham, ME) were driven along medial-lateral axis within the FEF recording chamber using a computer-controlled microelectrode drive (NAN Instruments; Israel.). Microstimulation pulse trains (100 msec, 300 Hz, 0.3 msec biphasic pulses, < 100 μA) were applied separately through each of these electrodes while the eye movements were observed. The location of an electrode was confirmed to be in the FEF if the microstimulation produced saccadic eye movements. The electrodes in medial FEF produced larger amplitude saccades than the ones in the lateral FEF, as has been reported previously (Bruce et al., 1985).

Neurons in ACC were recorded using arrays of one to three electrodes driven individually within the ACC recording chamber using a computer-controlled microelectrode drive (NAN Instruments; Israel). We attempted identifying neurons with direct projection or polysynaptic connectivity with FEF. In each experimental session, the activity of a single ACC neuron was isolated and monitored while single biphasic current pulses (0.15–0.3 ms per phase) were delivered to the FEF through one of the electrodes and a reference electrode in FEF chamber in contact with dura. Neurons were classified as antidromic if the stimulation elicited action potentials met certain criteria. These criteria include fixed latency given the axon has a fixed length, an action potential should be observed each time the stimulation pulse is delivered, and successful collision test (Lipski, 1981; Johnston and Everling, 2008) (see Fig. 1 D–F). For a collision test, we wait until the ACC neuron fires an action potential, and we simultaneously stimulate FEF. If the neuron sends a direct projection to FEF, the two action potentials (the endogenous ACC somal action potential and the elicited FEF action potential by stimulation) collide and will not be able to continue their path along the axon (given the absolute refractory period) and therefore, no action potential will be observed following the stimulation. The stimulation threshold for eliciting antidromic or orthodromic responses was defined as the current level that elicited an action potential ∼50% of the time. Threshold varied between 200 and 1500 μA. An orthodromic neuron was classified if it failed the collision test. Data collection initiated once an anti- or ortho-dromic (AO) neuron was identified. It should be mentioned that for Monkey 1, initial electrophysiological recordings sessions were conducted only recording the activity of ACC neurons without AO stimulation, but later added the AO stimulation to the recording sessions. In Monkey 2, the AO stimulation was performed from the beginning of the experiments. Action potential waveforms (40 kHz sampling frequency) were digitized, stored, and sorted off-line using two- and three-dimensional principal components analysis (Plexon Offline Sorter). In sessions when we could not identify an AO neuron, we resumed data collection without designating the isolated neurons as AO.

We also recorded multiunit activity in this study. We utilized a threshold of ± 3–5 standard deviations above or below the average raw signal to isolate the multiunit activity in ACC and FEF. In Offline Sorter, we inspected to ensure that multiunit activity displayed normal waveforms, and abnormal/artifactual waveform shapes were discarded. It should be mentioned that for Monkey 1, the ACC single unit recording experiments were conducted separately than the ACC and FEF multiunit recordings, whereas for Monkey 2, ACC single unit recordings, and ACC and FEF multiunit recordings were performed simultaneously during the same experimental sessions.

### Data analysis

Several task epochs were analyzed in further analysis. In the memory-guided saccade task, baseline epoch was the 500 ms prior to the presentation of the fixation point, fixation epoch was 50–250 ms following the fixation point onset, delay epoch was -1000–0 ms from offset of the central fixation point (which prompts the monkey to make a saccade towards the remembered target location), saccade epoch was -100–100 ms from saccade onset, post-saccadic epoch was 100–500 ms after saccade onset, and reward epoch was 50–450 ms after reward onset.

In the pro-/anti-saccade task, we designated the baseline epoch as the 500 ms prior to the presentation of the cue, preparatory epoch was -400–0 ms from stimulus onset, stimulus epoch was 50–250 ms following the stimulus onset, saccade epoch was -100–100 ms from saccade onset, post-saccadic epoch was 100–500 ms after saccade onset, and reward epoch was 50– 450 ms after reward onset.

Analysis of spiking activity was performed using a 3-way ANOVA with the factors: epoch (e.g., baseline vs. fixation epoch or baseline vs delay epoch), direction (of stimulus or saccade; e.g., 0° vs. 45° vs. 90°, etc.), and performance (correct vs. error trials). An additional factor, rule (e.g., prosaccade trial vs. antisaccade trial), was included in spiking analysis of the pro-/anti-saccade task (4-way ANOVA). Interaction between these factors indicated task-related differences in activity. The statistical significance for ANOVA tests was set at p < 0.05.

To bridge electrophysiological unit statistics with functional connectivity (FC), we reconstructed each neuron’s electrophysiological recording site onto MRI images. Recording sites are divided along the anteroposterior axis of ACC in 1 mm steps. After we determined the recording site for each neuron on the MR image, we averaged the FC of the recording site’s voxels with right medial and lateral FEF. In following analyses, we correlated (a) percentage of neurons with statistically significant epoch selectivity, direction preference, or performance related signal at each recording site with (b) average FC of the neurons’ recording site with right medial and/or lateral FEF (Fig. 6A).

We also investigated whether the connectivity of ACC recording sites with FEF results in facilitatory or inhibitory effects on the firing rate of the recorded neurons. To achieve this, we first calculated the “Activity Index” of each recorded neuron using the following formula:

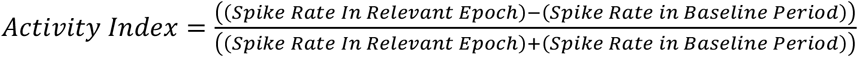

According to this formula, the Activity Index has a range of -1 to +1; if spike rate in the epoch is greater than baseline then Activity Index will be positive. If an epoch’s spike rate is less than baseline, Activity Index will be negative. We then compared the Activity Index of AO-identified neurons with that of all recorded neurons. Moreover, we correlated the average Activity Index of neurons recorded at each ACC recording site with the FC to FEF; only cells that had statistically significant difference between baseline and task epoch entered the correlation analysis.

Murphy and colleagues have suggested that multiunit activity is better correlated with FC than other electrophysiological correlates, such as calcium imaging and banded LFP power (Murphy et al., 2018). To further examine the relationship between FC and electrophysiological correlates, we investigated whether the correlated multiunit activity between ACC and FEF is related to FC. We first calculated the correlation of the multiunit activity in each ACC recording site, with multiunit activity of the medial and lateral FEF across all task epochs and identified the multiunit channel pairs with statistically significant correlation (either positive or negative; Pearson’s correlation, refer to Fig. 7E for an example of a positive multiunit pair and Fig. 7F for an example of a negative multiunit pair). We computed the proportion of significantly correlated multiunit pairs for each ACC recording site (medial and lateral FEF separately). Across recording sites, we tested for correlation between (a) the proportion of significant multiunit pairs and (b) rs-fMRI FC with FEF (both positive and negative FC, separately).

## Results

Two adult rhesus macaques performed the memory-guided saccade task and the pro-/anti-saccade task during electrophysiological recordings in ACC (Fig. 1). In the memory-guided saccade task, the correct performance rate was 94.68% for Monkey 1 and 89.97% for Monkey 2. In the pro-saccade task, correct performance rate was 91.79% for Monkey 1 and 93.13% for Monkey 2. In anti-saccade trials, correct performance rate was 86.74% for Monkey 1, and 86.15% for Monkey 2.

### Single unit spiking activity in memory-guided saccade task

We first attempted to characterize the activity of the single neurons recorded in the ACC. Figure 2A shows the percentage of neurons with significant activity in different epochs of the memory-guided saccade task. From the original 425 neurons, some were removed if a minimum threshold of 40 performed trials was not reached, resulting in 381 neurons for further analysis. It should be mentioned that only 24 out of 381 total analyzed neurons were identified as AO-identified neurons and therefore these results should be interpreted cautiously due to low number of cells. Given monkeys’ performance was ∼95% in the memory-guided saccade task, we did not have enough error trials and could not measure performance-related signal during memory-guided saccade tasks. We observed the highest percentage of epoch-selectivity spiking activity in reward (21.67%) and post-saccadic (20.88%) epochs (Fig. 2A). Percentage of neurons with direction-selective spiking activity was above chance level in visual (6.52%), delay (7.83%), post-saccadic (9.39%), and reward (9.13%) epochs using binomial distribution at p <

**Figure 2.**
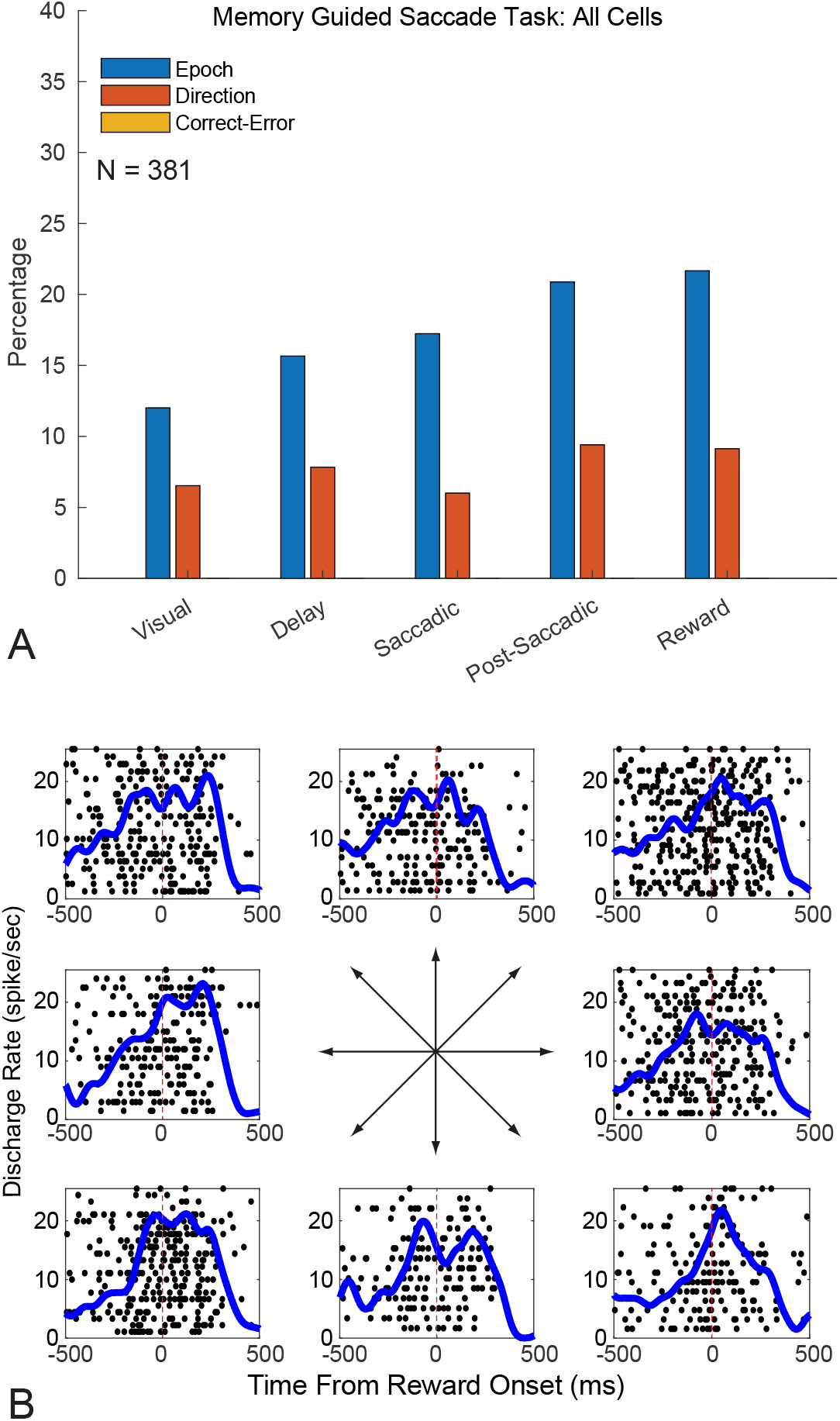
Neuronal selectivity to the memory-guided saccade task. (A) Percentage of significant epoch-, direction-, and performance-selective neurons across different epochs of the memory-guided saccade task. No neurons had significant performance-related activity, due to the low number of error trials in memory-guided saccade task. (B) Raster plots and peri-stimulus time histograms of an example neuron with reward-related activity. Each dot represents an action potential. The activity of the neuron is depicted separately for each of the 8 target locations.

0.05 threshold (Fig. 2A). The percentage of epoch-selective neurons was significantly higher than the percentage of direction-selective neurons. Figure 2B shows an example neuron with reward-related activity.

### Single unit spiking activity in pro-/anti-saccade task

Next, we explored the percentage of neurons with selective activity across different epochs during the pro-/anti-saccade task. Similar to the memory-guided saccade task, the highest percentage was epoch selectivity in post-saccadic (28.24%) and reward (27.11%) epochs (Fig. 3A). We also explored the percentage of neurons that differentiated between pro- and anti-saccades across different task epochs (“rule-selective”). Percentage of rule-selective cells was lower than epoch-selective cells (Fig. 3A). Interestingly, the percentage of neurons with performance selectivity (differentiating between correct and error trials) were higher in post-saccadic and reward epochs (compared to preparatory, visual, and saccadic epochs), reflective of the performance-monitoring role of ACC neurons. Figure 3B shows an example neuron with significant post-saccadic activity in the pro-/anti-saccade task.

**Figure 3.**
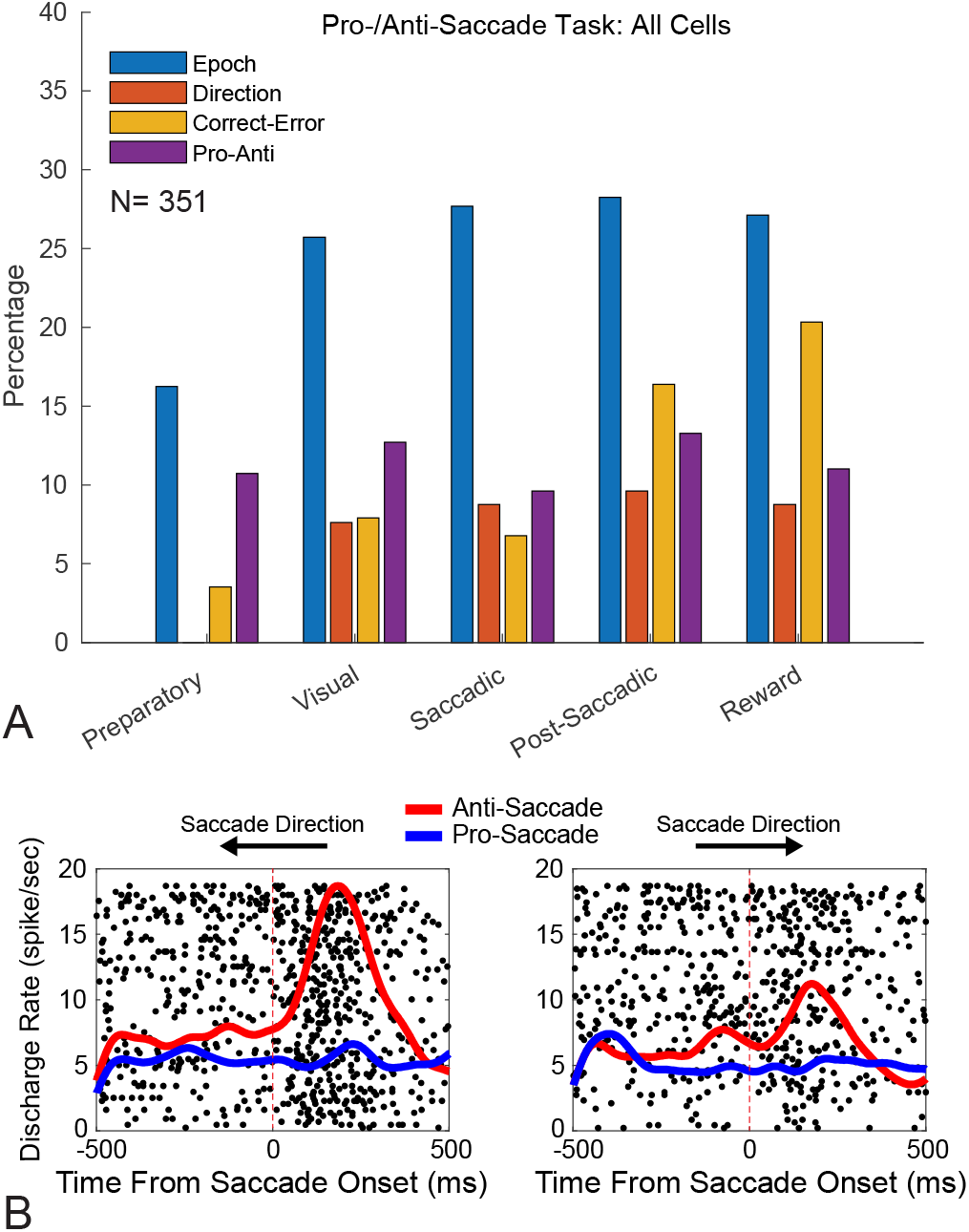
Neuronal selectivity to the pro- and anti-saccade task. (A) Percentage of significant epoch-, direction-, performance-, and rule-selective neurons across different epochs of the pro- and anti-saccade task. (B) Raster plots and peri-stimulus time histograms of an example neuron with epoch-, direction- and rule-selectivity in the post-saccadic epoch. Each dot represents an action potential. The activity of the neuron is depicted for two saccade directions.

### Anti- and ortho-dromically (AO) identified neurons: ACC neurons projecting to FEF

We explored the activity of AO-identified neurons in different task epochs in both the memory-guided saccade task and the pro-/anti-saccade task (Fig. 4A and 4B, respectively). Overall, we observed similar patterns of selectivity for the AO-identified neurons as for all recorded neurons: epoch-selective neurons were more prevalent than direction-selective or performance-selective neurons. The only exception was that we observed similar number of epoch-selective and direction-selective neurons in the saccade epoch of memory-guided saccade task—both exceeded the chance level utilizing binomial distribution at p < 0.05 level.

**Figure 4.**
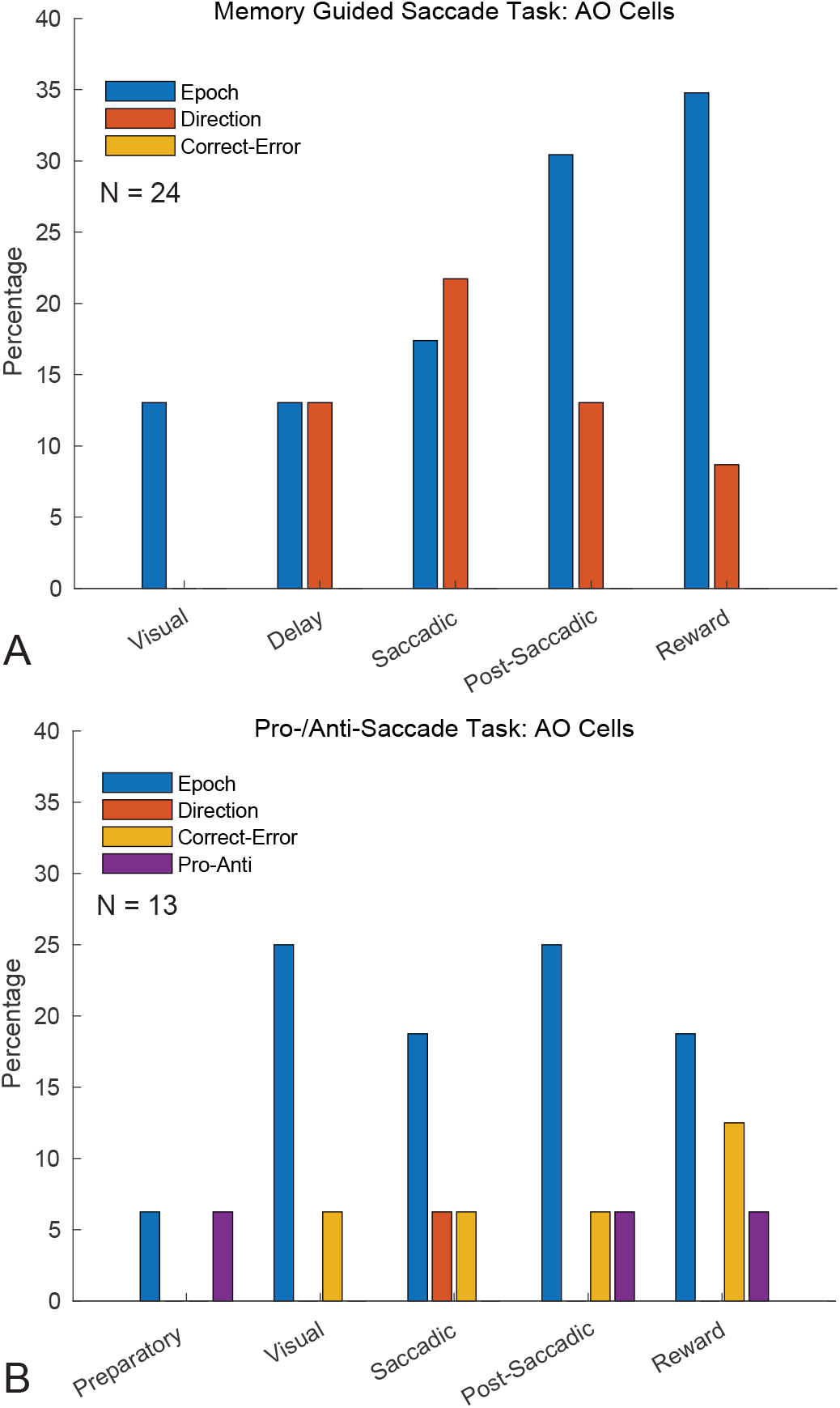
Task selectivity of anti- and ortho-dromic (AO) neurons (A) Percentage of significant epoch-, direction-, and performance-selective AO-identified neurons across different epochs of the memory-guided saccade task. (B) Percentage of significant epoch-, direction-, performance-, and rule-selective AO-identified neurons across different epochs of the pro- and anti-saccade task.

### Connectivity of AO-identified ACC neurons with FEF

In our previous rs-fMRI study, we demonstrated that medial FEF, but not lateral FEF, show strong functional rs-fMRI connectivity (Babapoor-Farrokhran et al., 2013). Here, we combined rs-fMRI (Fig. 5) and electrophysiology of AO-identified neurons to test ACC connectivity with medial, or lateral FEF. Figure 5 shows the FEF seed regions and resultant FC maps (both positive and negative). ACC regions, corresponding to electrophysiological recording sites, are shown for each monkey.

**Figure 5.**
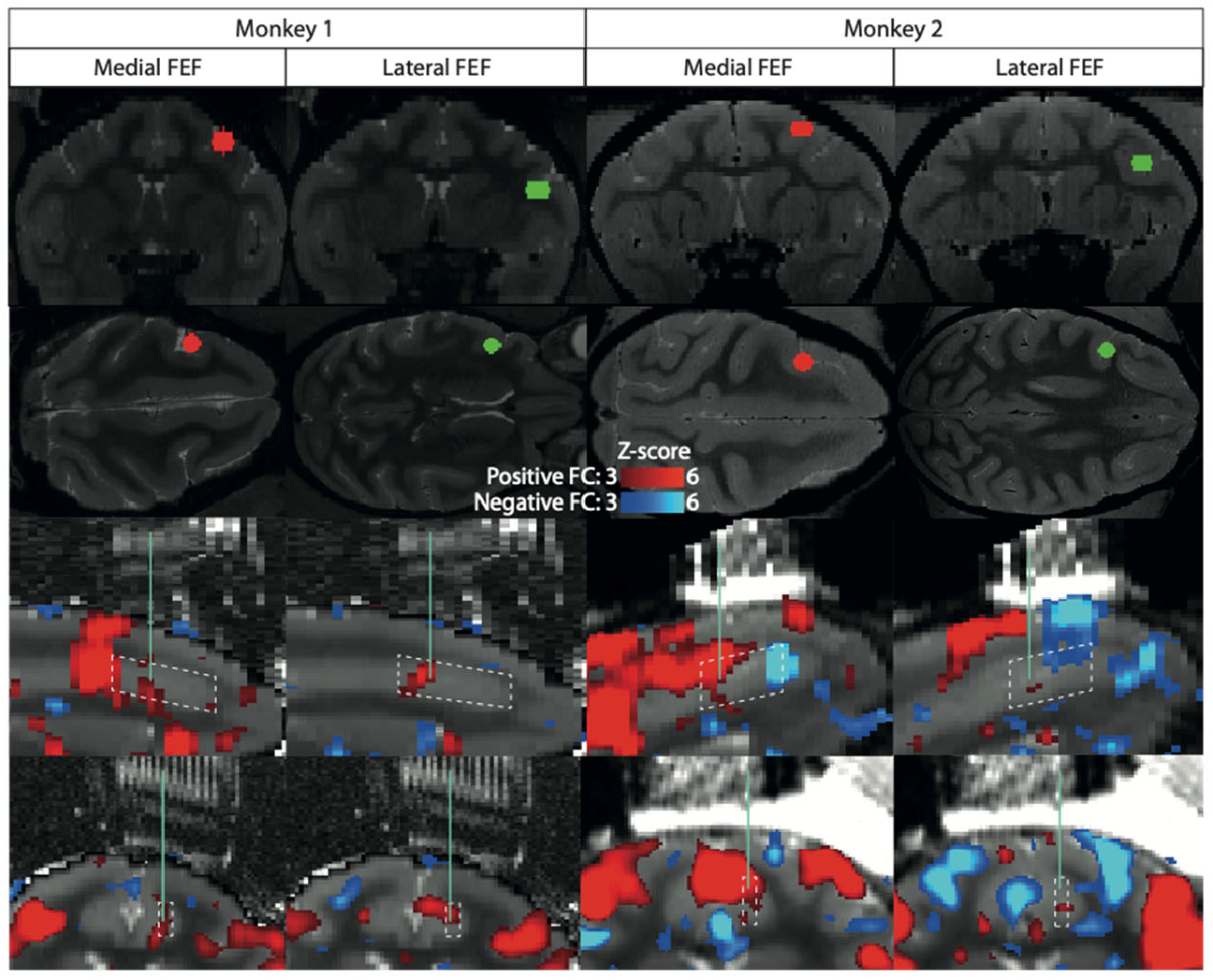
Functional connectivity (FC) of ACC and FEF. (Top) Placement of seed regions in medial (red) and lateral (green) FEF in Monkey 1 (Left) and Monkey 2 (Right); first row is coronal, and second row is transverse MRI sections. (Bottom) Functional connectivity of medial and lateral FEF. Third row is sagittal and forth row is coronal section. The positively correlated areas are shown in red and negatively correlated areas are shown in blue. The green lines on the bottom two rows illustrate the trajectory of a recording electrode. The recording sites in each monkey are designated by dotted white lines.

We found that most of the orthodromic identified ACC neurons were located in the medial FEF (Fig. 6A, 18/25 cells, p < 0.0001 binomial test), which is in line with our previously reported rs-fMRI results. Moreover, both of the antidromically identified neurons were connected with medial FEF. We found that the AO-identified neurons were recorded from sites with higher trend of FC z-scores (Fig. 6B,C); however this did not reach statistical significance due to the low number of AO sites (n = 6 AO sites, n = 24 total sites, mean of AO sites’ medial FEF positive FC z-score = 2.96, mean of all sites’ medial FEF positive FC z-score = 2.0167, Kruskal-Wallis p-value =0.0969). Generally, medial FEF had greater positive FC z-scores (mean all sites = 2.0167; mean AO sites = 2.96) compared to lateral FEF (mean all sites = 0.9941, mean AO sites= 1.4431, all sites’ Kruskal-Wallis p = 0.0044, AO sites’ Kruskal-Wallis p = 0.0547). With respect to negative FC z-score, medial FEF had significantly greater FC z-scores (mean all sites = -0.5503, mean AO sites = -0.0303) compared to lateral FEF (mean all sites = -0.7951, mean AO sites = -0.5197, all sites’ Kruskal-Wallis p = 0.0011, AO sites’ Kruskal-Wallis p = 0.0163).

**Figure 6.**
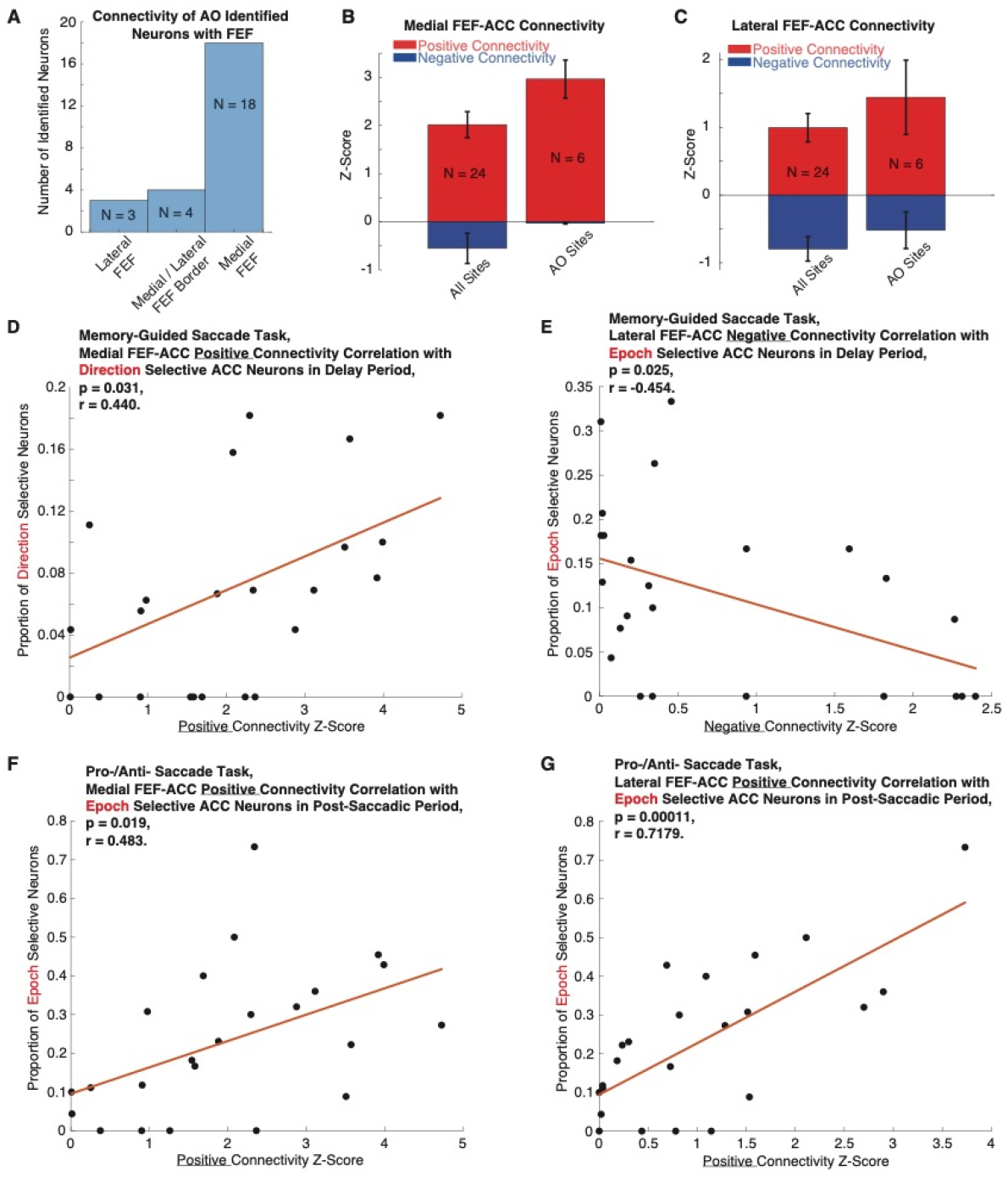
Interaction between the imaging and electrophysiological connectivity of FEF and ACC. (A) Histogram of the number of AO-identified ACC neurons with medial, lateral, medial/lateral bordering areas of FEF. (B) Functional connectivity (FC) of all cells compared to the FC of AO-identified cells with medial FEF (mean +/-standard error of the mean). Positive correlation is shown in red and negative correlation is shown in blue. (C) Same as B but for lateral FEF. (D) Correlation of medial FEF-ACC positive FC with proportion of the direction-selective cells in the delay epoch of the memory-guided saccade task. (E) Correlation of lateral FEF-ACC negative FC with proportion of the epoch-selective cells in the delay epoch of the memory-guided saccade task. (F) Correlation of medial FEF-ACC positive FC with proportion of the epoch-selective cells in the post-saccadic epoch of the pro- and anti-saccade task. (G) Correlation of lateral FEF-ACC positive FC with proportion of the epoch-selective cells in the post-saccadic epoch of the pro- and anti-saccade task.

### Correlation of ACC spiking activity with connectivity to FEF

We also examined whether there is a correlation between (a) percentage of epoch-, direction-, or performance-selective neurons within each ACC recording site with (b) the functional connectivity (FC) of the recording site with medial and lateral FEF. These correlations were determined across different task epochs. Both positive and negative FC were examined.

In the MGS task, we found significant correlation between medial FEF positive connectivity z-score and percentage of neurons with epoch selectivity in the post-saccadic epoch (Pearson’s correlation, r = 0.459, p = 0.023), and direction selectivity in delay (r = 0.440, p = 0.031; Fig. 6D) and saccadic (r = 0.406, p = 0.0488) epochs. That is, ACC sites with higher proportions of epoch- or direction-selective neurons often had greater FC with medial FEF. However, there was no correlation between medial FEF *negative* connectivity z-score and percentage of ACC neurons with epoch selectivity.

Next, we analyzed lateral FEF during MGS performance. There was no correlation between lateral FEF *positive* connectivity z-score and percentage of neurons with task selectivity in MGS. However, there was a significant negative correlation between *negative* connectivity z-score and percentage of neurons with delay epoch selectivity (r = -0.454, p = 0.0257; Fig. 6E) and direction selectivity in the delay epoch (r = -0.473, p = 0.0195). Said differently, ACC sites with higher proportions of epoch- or direction-selective neurons during the delay epoch often had greater *negative* FC with lateral FEF.

In the pro-/anti-saccade task, there was a significant correlation between medial FEF *positive* connectivity z-score and percentage of neurons with epoch selectivity in visual (r = 0.481, p = 0.02) and post-saccadic (r = 0.483, p = 0.019; Fig. 6F) epochs. There was no correlation between medial FEF *negative* connectivity z-score and percentage of neurons with epoch selectivity.

In lateral FEF during pro-/anti-saccade task, *positive* connectivity z-score significantly correlated with percentage of neurons with epoch selectivity during all task epochs: visual epoch (r = 0.698, p = 0.0002), preparatory epoch (r = 0.613, p = 0.0018), saccadic epoch (r = 0.685, p = 0.0003), post-saccadic epoch (r = 0.718, p = 0.00011; Fig. 6G), and reward epoch (r = 0.528, p = 0.009). In addition, there was a significant negative correlation between lateral FEF negative connectivity z-score and percentage of neurons with epoch selectivity in visual (r = -0.507, p = 0.013), and post-saccadic (r = -0.431, p = 0.039) epochs.

### Relationship of spike rate facilitation or inhibition with connectivity to FEF

We used Activity Index measure to investigate whether the connectivity of the ACC recording sites with FEF influences the spike rate of neurons during the task epochs. We did not observe any significant difference between the Activity Index of AO-identified cells with that of the population of cells in any epoch of MGS or pro-/anti-saccade task (t-test, p > 0.05, Fig. 7A & B). We also explored the correlation of the average Activity Index in each ACC recording site with ACC-FEF FC. The only statistically significant correlation was between ACC Activity Index in the MGS reward epoch and positive FC with lateral FEF (Pearson’s correlation, r = 0.463, p = 0.022, Fig. 7C); however, this significance did not survive the Bonferroni correction for multiple comparisons. Across all other MGS epochs and the pro-/anti-saccade task, there was no correlation between the ACC Activity Index with positive or negative FC of medial or lateral FEF (e.g., Fig. 7D).

**Figure 7.**
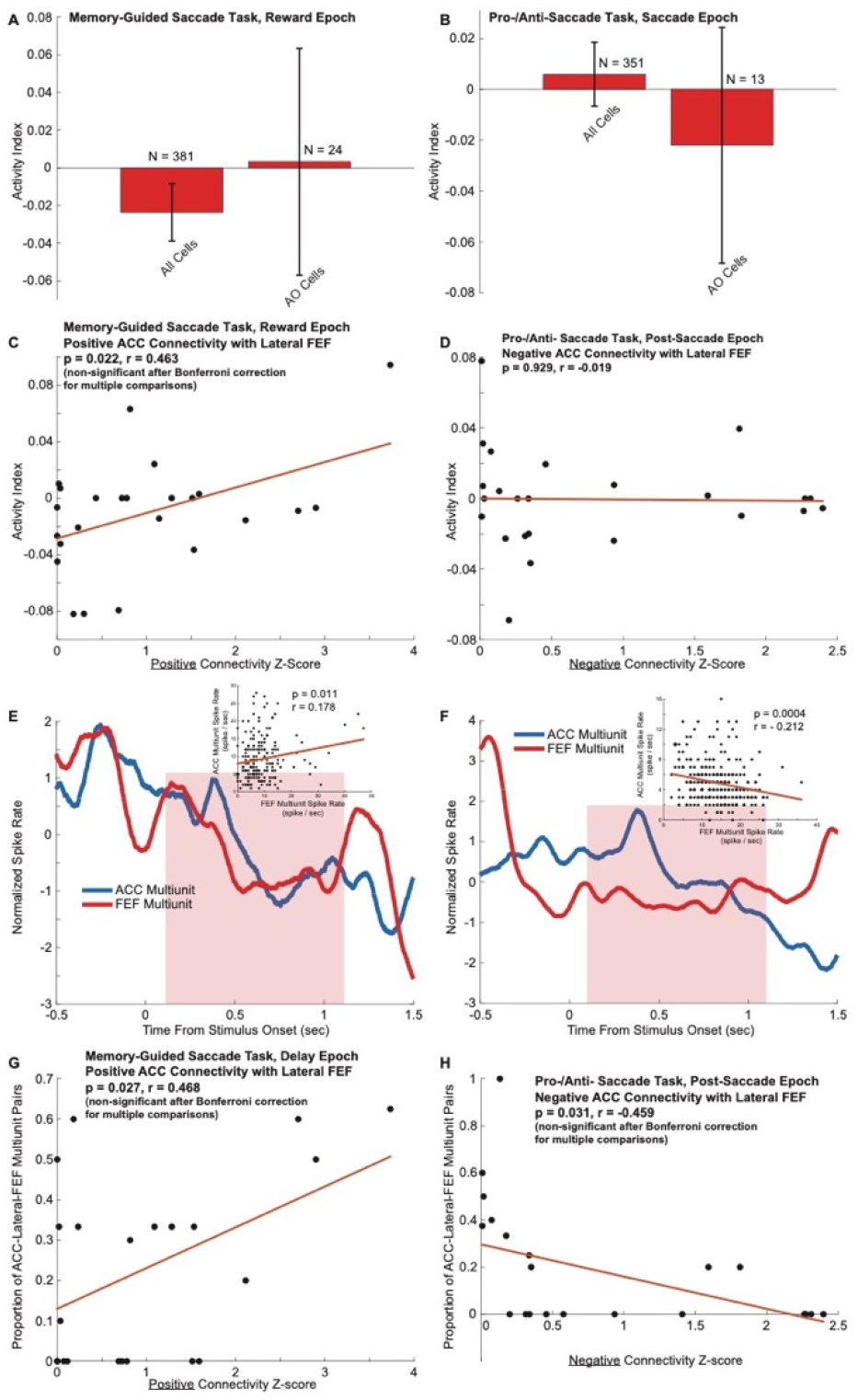
Relationship between task-related spiking and ACC-FEF connectivity. (A) Activity Index during the reward epoch of the memory-guided saccade task for all cells and AO-identified cells (mean +/-standard error of the mean). There was no significant difference between all cells and AO-identified cells (t-test, p > 0.05). (B) No significant difference in Activity Index during the saccade epoch of the pro-/anti-saccade task between all cells and AO-identified cells (t-test, p > 0.05). (C) Correlation of lateral FEF-ACC positive FC with Activity Index during reward epoch of memory-guided saccade. (D) Correlation of lateral FEF-ACC negative FC with Activity Index during post-saccade epoch of pro-/anti-saccade task. (E) Example of an ACC and lateral FEF multiunit pair with *positive* correlation of their activity (“positive” multiunit pair) in the delay epoch of the MGS. The shaded area delineates the delay epoch. Scatter plot of multiunit activity for each region and delay epoch is overlaid on the figure. (F) Example of an ACC and lateral FEF multiunit pair with *negative* correlation of their activity (“negative” multiunit pairs) in the delay period of the MGS. (G) Across recording sites, proportion of significant ACC-lateral-FEF multiunit pairs is correlated with positive FC in the delay epoch of the memory-guided saccade task. (H) Across recording sites, proportion of significant ACC-lateral-FEF multiunit pairs is correlated with negative FC in the post-saccadic epoch of the pro-/anti-saccade task.

### Relationship between ACC-FEF multiunit correlation and ACC-FEF functional connectivity

We examined whether correlations between ACC and FEF multiunit activity was related to the ACC-FEF FC. Over the course of different trial epochs, was multiunit activity from ACC and FEF significantly correlated (e.g., Fig. 7E,F)? We observed a positive correlation between the proportion of significant ACC-lateral-FEF multiunit pairs and positive ACC-lateral-FEF FC in the delay epoch of the MGS task (Pearson’s correlation, r = 0.468, p = 0.027, Fig. 7G). That is, ACC sites with a greater proportion of multiunit pairs led to stronger positive FC with lateral FEF. We also found a negative correlation between the proportion of significant ACC-lateral-FEF multiunit pairs and negative ACC-lateral-FEF FC (or anti-correlations) in the post-saccadic epoch of the pro-/anti-saccade task (r = -0.459, p = 0.031, Fig. 7H). That is, ACC sites with a greater proportion of multiunit pairs led to reduced negative FC. It should be emphasized that although these results were statistically significant, they did not reach the statistical significance threshold when correction for multiple comparisons was applied (Bonferroni correction for multiple comparisons). We also examined multiunit pairs between ACC and medial FEF, but found no correlation between proportion of significant multiunit pair and FC.

For each pair, we then examined whether ACC and FEF multiunit activities were positively correlated or negatively correlated. Were ACC and FEF multiunit activity moving together or in opposing directions during the course of each trial? There was not a significant relationship for one direction or the other in MGS and a weak relationship in pro-/anti-saccade task. Out of 27 ACC-lateral-FEF multiunit pairs with significantly correlated activity in the delay epoch of MGS, 11 were negatively correlated and 16 were positively correlated (Binomial test, p > 0.05). During the post-saccadic epoch of pro-/anti-saccade task, 7 of 22 ACC-lateral-FEF multiunit pairs were negatively correlated and 15 were positively correlated (p = 0.0407). During this task epoch, multiunit pairs were more likely to change their activity in a coherent fashion. This is interesting, given the proportion of these multiunit pairs during pro-/anti-saccade task had negative correlation with negative FC (Fig. 7H).

To further explore the neurophysiological underpinnings of FC, we examined whether positive multiunit correlation was indicative of positive FC, or whether negative multiunit correlation was indicative of negative FC, or vice versa. Within “positive” multiunit pairs (i.e., ACC multiunit activity and lateral FEF multiunit activity were positively correlated, Fig. 7E), we did not find a statistically significant correlation between proportion of significant pairs at each site and positive FC. Similarly, “positive” multiunit pair proportion was not correlated with negative FC. An unexpected pattern was found for “negative” multiunit pairs (Fig. 7F): proportion was positively correlated with positive lateral FEF FC during delay epoch of MGS (r = 0.657, p = 0.00088) and during post-saccadic epoch of pro-/anti-saccade task (r = 0.587, p = 0.0048).

Similarly, “negative” multiunit pairs proportion was negatively correlated with negative FC (post-saccade epoch of pro-/anti-saccade task; r = -0.473, p = 0.0258). Thus, “negative” multiunit pairs (wherein ACC and lateral FEF multiunit activity were negatively correlated; e.g., Fig. 7F) were more likely to be associated with significant FC than “positive” multiunit pairs. This relationship was inverted for positive FC and negative FC. That is, greater number of significant “negative” multiunit pairs corresponds with greater positive FC and reduced magnitude of negative FC. We also performed Bonferroni correction for multiple comparisons. The only relationship that survived the Bonferroni correction was between the proportion of “negative” multiunit pairs and positive lateral FEF FC, which was positively correlated during delay epoch of MGS.

## Discussion

This investigation evaluated the functional properties of ACC in saccade tasks and identified ACC neurons that received input from or projected to the FEF. In addition, we compared the spiking activity of FEF-connected ACC neurons to FC, as determined by rs-fMRI. Earlier work had suggested that ACC has only weak connections to FEF, and instead supplementary eye fields (SEF) are better-positioned to directly influence saccade production (Schall et al., 1993, 2002). However, more recent imaging evidence and the current study support a direct ACC-to-FEF connection. By combining electrophysiology and imaging techniques, we benefit from the strengths of each method. Single-unit electrophysiology provides the high informational content and high temporal resolution during relevant cognitive behavior whereas rs-fMRI provides wide spatial coverage, allowing comparisons between different subregions of ACC and FEF.

By placing an FEF seed, rs-fMRI shows FC between ACC and FEF, particularly in medial regions of FEF and posterior segments of ACC (Fig. 5). Previous imaging work has shown medial vs lateral FEF form separate brain-wide connections, with medial FEF connecting to regions related to reaching and cognitive control, whereas lateral FEF connects with regions related to object manipulation and fixation of gaze (Babapoor-Farrokhran et al., 2013). A greater connection to medial FEF is consistent with these regions’ shared involvement in goal-directed action and cognitive control. This FEF-connected region of ACC is likely overlapping with “cingulate eye fields” established in previous work using retrograde tracers in FEF (Wang et al., 2004). (Vezoli et al., 2021) recently reported that rs-fMRI FC is correlated with LFP coherence and granger causality (especially feedforward gamma activity), further evidence of a direct and functional connection between these regions. Connectivity between these regions is consistent with anatomy (Barbas and Mesulam, 1981; Huerta et al., 1987; Bates and Goldman-Rakic, 1993; Stanton et al., 1993, 1995; Wang et al., 2004), and previous FC (Hutchison et al., 2012).

Previous studies have also examined local field potential (LFP) coherence between these prefrontal regions. LFP of these regions are coherent in the theta and low beta frequency bands (Womelsdorf et al., 2014; Babapoor-Farrokhran et al., 2017). Specifically, theta coherence was bidirectional between ACC and FEF, whereas beta connectivity preferred the ACC-to-FEF direction (as determined with Granger causality) during the working memory delay epoch. This is consistent with the proposed roles of theta in long-range binding of cortical regions and attentional sampling (Fries, 2015; Fiebelkorn et al., 2018) and beta oscillations in top-down control (Bastos et al., 2015, 2020).

Analysis of all ACC neurons in the MGS and anti-saccade task shows that units are responsive to specific task epochs (e.g., visual stimulus, delay epoch, saccadic response, reward epoch) and responsive to specific saccade directions (Fig. 2A). This supports the role of ACC in saccadic response and processing of specific task timing, events, and stimuli. During the anti-saccade task, some neurons were additionally selective for specific task rules—pro-saccade or anti-saccade. Representation of abstract rules is consistent with ACC’s involvement in cognitive control, and in the case of anti-saccades, ACC’s involvement in response inhibition.

Here, ACC sites were delineated based on electrophysiological recording locations, such that FC could be compared with electrophysiological characteristics. Previous work with imaging alone had detailed the reciprocal connections between these two regions of frontal cortex, but recording during cognitive behavior in this study permits functional testing of the hypothesized role of ACC in FEF saccade generation and inhibition. Consistent with our hypothesis, ACC sites with AO-connected neurons had a trend of greater functional connectivity scores compared to non-connected sites and this was true for both medial and lateral FEF, although this did not reach statistical significance due to low number of the AO-identified cells (Fig. 6B,C). Orthodromically-connected ACC neurons at these sites are evidence of interaction between ACC and FEF, either directly (in the case of antidromically-identified neurons) or indirectly (orthodromic connectivity can result from two or three synapses, e.g., through thalamus). It is noteworthy that this interaction is bidirectional: ACC can provide top-down influence to FEF and FEF provides bottom-up influence over ACC (Babapoor-Farrokhran et al., 2017). Both FEF and ACC are part of the dorsal attention network and cognitive control, but do they interact during cognitive saccadic behavior?

### ACC-FEF interaction in motor preparation and saccade generation

Consistent with early ACC work describing directional delay activity (Niki and Watanabe, 1979), some FEF-connected ACC neurons were specifically active during the motor preparation period (delay epoch) of memory guided saccades (Fig. 4A). Previous imaging studies support that ACC influences FEF during presentation of visual stimuli (Ding et al., 2009) and immediate motor planning to a target (Schmid et al., 2001). However, it is noteworthy that delay was not the most prominent task epoch for cell selectivity, and relationship between task-selective neurons and functional connectivity was not as strong during the delay (e.g., compared to saccadic epochs). If ACC indeed exerts cognitive control over FEF, perhaps this input is more likely to occur during the saccade epochs. For example, the greatest number of epoch-selective neurons were found during saccadic, post-saccadic, and reward task period. This indicates ACC may exert greatest control during saccade generation and afterwards. Considering the high proportion of direction-selective ACC neurons in these saccadic epochs, this appears to exemplify the dorsal attention network and influence from ACC onto FEF to generate saccadic outputs in specific directions, and possibly transmission of direction-related information from FEF to the ACC during these epochs. A role of ACC in saccade generation is supported by earlier work, showing that microstimulation of dorsal ACC can evoke saccades at < 60 μA (Mitz and Godschalk, 1989).

### ACC-FEF interaction in reward and conflict monitoring

Pierrot-Deseilligny et al., (2002) proposes ACC adds a motivational salience, perhaps enhancing preparation of intentional saccades, and nudging the circuits towards saccade threshold with task-relevant timing. This added salience may be related to the fact that compared to other prefrontal regions, ACC is strongly connected with the limbic and reward centers, including orbitofrontal cortex (Tang et al., 2019), amygdala (Vogt, 2005), and ventral tegmental area (via striatum, (Haber and Knutson, 2010)). ACC is therefore well-positioned to evaluate rewards or unexpected omission of reward (Niki and Watanabe, 1979) as a result from goal-directed saccade. Aptly, here we report many reward-epoch-selective ACC neurons.

Task-related neurons were more likely to be found in the later task epochs, such as post-saccadic and reward periods. This finding coincides with the well-known role of ACC in conflict monitoring (Johnston et al., 2007; Womelsdorf et al., 2010). As part of the executive control network, ACC is proposed to be active during uncertainty (Fan et al., 2014) and resolve conflicts between competing processes (Xuan et al., 2016). Imaging studies have described ACC activation when subjects must inhibit competing responses (Posner and DiGirolamo, 1998) and event-related potentials after errors appear to originate from ACC (Dehaene et al., 1994).

### ACC-FEF interaction in cognitive control and response inhibition

Many neurons in the full spiking dataset were selective for abstract rules. Spike rate variability in these units were explained by preference for either pro-saccade rule or anti-saccade rule, suggesting they were involved in these task conditions. The anti-saccade task is often used as a proxy for cognitive control, such as inhibition of prepotent responses (Johnston et al., 2007), and performance is disrupted across several conditions linked to disrupted cognitive behavior, including Alzheimer’s disease, Parkinson’s disease, and schizophrenia (Fukushima et al., 1994; Kaufman et al., 2010). The presence of anti-saccade selective neurons in ACC, shown here, may support its role in inhibiting erroneous prosaccades during anti-saccade trials. Although lesion studies often implicate dorsolateral prefrontal region in suppressing saccades (Pierrot-Deseilligny et al., 1991, 2002), similar lesion studies have reported anti-saccade errors after lesions to ACC (Gaymard et al., 1998; Milea et al., 2003). Thus, in addition to dorsolateral PFC, ACC may additionally suppress saccade generation motifs in FEF. Imaging studies have shown greater activation in ACC during anti-saccades than prosaccades (Paus et al., 1993; Doricchi et al., 1997; Matsuda et al., 2004; Ford et al., 2005; Manoach et al., 2007; Brown et al., 2008), further implicating a role in cognitive control of response inhibition. Curiously, a relationship between proportion of anti-saccade selective neurons and functional connectivity strength was not significant across ACC sites, but such a relationship was prevalent for saccade-epoch and direction-selective neurons. This may represent a difference in specialization between ACC subregions, with this subregion having a greater representation for directional saccade generation (e.g., cingulate eye fields, (Wang et al., 2004)) as opposed to response suppression, and further support the notion that ACC might be receiving directional information from FEF in these areas.

### Positive vs. negative functional connectivity

Positive rs-fMRI functional connectivity correlations are thought to reflect monosynaptic or polysynaptic anatomical connections (Babapoor-Farrokhran et al., 2013). However, the significance of negative rs-fMRI correlations (also known as anti-correlations) is still debated. It has been previously shown that global signal removal during rs-fMRI pre-processing could introduce artifactual anti-correlations in the results (Murphy et al., 2009). However, others have argued that anti-correlated brain networks cannot be explained solely by global signal regression during preprocessing and these negative correlations instead represent physiologic brain networks (Fox et al., 2009). For example, the default mode network is known to be anti-correlated with the dorsal attention network (Fox et al., 2005, 2009). Rs-fMRI anticorrelations are proposed to result from polysynaptic connections and transition time delays between networks (Greicius et al., 2003; Deco et al., 2009). Further, anti-correlated rs-fMRI has now garnered attention in clinical trials. In recent studies for treatment-resistant depression, the most beneficial effect was found after rTMS to an individual-specific region of dorsolateral PFC that was the most anticorrelated with subgenual ACC (Fox et al., 2012, 2014; Cole et al., 2020).

To the best of our knowledge, our study is the first to test the electrophysiological underpinnings of positive vs negative FC in vivo by using the anti-dromic stimulation technique—considered the gold standard of electrophysiological connectivity analysis in vivo. We observed that a statistically significant majority of the AO-identified neurons were connected to the medial FEF, which has positive FC with ACC. Conversely, fewer AO-identified neurons were connected to lateral FEF, which has mostly negative FC to ACC. Together, this suggests anatomical connectivity (anti-/ortho-dromically verified mono-/polysynaptic connections) is related to positive FC but not negative FC. Further, a lack of mono-/polysynaptic anatomical connectivity may be related to negative FC, but more experiments are required to confirm this proposal. Medial and lateral FEF have previously been shown to have different anatomical and network characteristics (Preuss and Goldman-Rakic, 1991). We have previously shown that medial FEF has stronger rs-fMRI functional connectivity to ACC compared to the lateral FEF (Babapoor-Farrokhran et al., 2013). This contrast of medial vs lateral FEF is interesting given the current results, which identify ACC neurons as having more AO-identified neurons connected with medial FEF. Although our findings do not completely rule out the presence of a physiologic underpinning for anti-correlated networks, these results support that positive FC correlation are most likely driven by anti- or ortho-dromically defined anatomical connections. In this region of ACC, anticorrelated brain regions appear to have fewer AO-identified connections and less anatomical connectivity.

We also explored whether anti-/ortho-dromic connectivity or rs-fMRI connectivity could reflect possible excitatory or inhibitory effect on the activity of neurons. We utilized Activity Index, which shows increased or decreased activity of the neurons compared to the baseline period. The only statistically significant finding was a positive correlation between ACC Activity Index in the MGS reward epoch and positive FC with lateral FEF and even this result was not significant after Bonferroni correction for multiple comparisons. There was no other correlation between the Activity Index and positive or negative FC with medial or lateral FEF across other task epochs. A lack of relationship between Activity Index and positive/negative FC may support the proposal that FC is not related to excitation or suppression of neuronal activity. Furthermore, we investigated whether multiunit activity correlation could explain FC variability across different recording sites. We found a higher proportion of significantly correlated ACC-lateral-FEF multiunit pairs in ACC sites with greater positive FC to lateral FEF (delay epoch of MGS task). In contrast, sites with higher proportion of significant multiunit pairs had a reduced magnitude of negative FC (post-saccadic epoch of pro-/anti-saccade task; Fig. 7G, H). However, all but one of these effects were not significant after tests for multiple comparisons. Interestingly, the type of relationship between ACC and lateral FEF multiunit activity was associated with strength of FC. ACC-lateral-FEF multiunit pairs were considered either “positive” or “negative” (i.e., ACC and FEF multiunit activities were positively or negative correlated – varying together or in opposing directions). Within “negative” ACC-lateral-FEF multiunit pairs (ACC and FEF multiunit activity were negatively correlated), the proportion of pairs at each recording site was positively correlated with positive FC and negatively correlated with negative FC. This may indicate that “negative” multiunit relationships between ACC and FEF are more indicative of a functional connection and therefore mono- or poly-synaptic anatomical connections. It is presently unclear whether “negative” multiunit relationships represent inhibitory relationships between brain regions. However, since multiunit activity represents average spiking of neurons in the local vicinity of a recording electrode, one may speculate that a “negative” relationship between regions indicates a mutually or unidirectionally suppressive relationship.

Overall, our electrophysiological evidence supports that positive FC reflects anatomical connectivity (mono- or poly-synaptic), but the same cannot be said for negative FC (anticorrelations). We observed that the signs of the correlation between task selectivity and positive vs negative functional connectivity are opposite. There is often a positive correlation of task selectivity with positive functional connectivity. In contrast, there is often a negative correlation of task selectivity with negative functional connectivity. Furthermore, we found suggestive evidence that the ACC-FEF multiunit coherence was more prevalent in areas with positive FC and less prevalent in areas with negative FC. It has been suggested that negative FC (anticorrelations) either have neural or vascular origin (Liu et al., 2015). Given that we found weak or no evidence of neuronal signals driving the negative FC (anticorrelations), it is possible that negative FC networks are instead driven by vascular supply. Blood supply in the brain is limited. When more blood is directed to an engaged brain area, inevitably other areas will receive less vascular supply. Therefore, these anticorrelations may be the result of blood flow dynamics (Liu et al., 2015), possibly acting on longer timescales (seconds), as opposed to fast mono-/poly-synaptic connections (on the timescale of tens to hundreds of milliseconds). Our results support this, since we do not find a relationship between mono-/poly-synaptic connections (AO-identified cells) and negative FC. Needless to say, further research is required to better elucidate the mechanisms of rs-fMRI correlations and anticorrelations.

By leveraging anti- and ortho-dromic stimulation, this investigation provided evidence of mono- or polysynaptic connections between ACC and FEF during cognitive saccade tasks. These ACC recording sites were found to have the greatest functional connectivity with medial subregions of FEF, which are related to cognitive control (compared to lateral FEF). A limitation of this study is the relatively small region of ACC that was sampled compared to the large volume of ACC and its many specialized subregions. Although the subregion sampled with electrophysiology contained a large representation for saccade- and direction-selective neurons that likely corresponding to cingulate eye fields, different subregions (slightly more posterior) may have greater cognitive control properties, such as influencing inhibition of saccades in FEF. Additionally, our results yield insights towards to electrophysiological underpinning of negative functional connectivity. Future studies utilizing high-density multi-contact arrays may provide valuable insight on the different subregions of ACC and their interactions with FEF.

